# STIM1 interacts with HCN4 channels to coordinate diastolic depolarization in the mouse Sinoatrial node

**DOI:** 10.1101/2023.05.03.539287

**Authors:** Hengtao Zhang, Victoria Graham, Igor Nepliouev, Jonathan A. Stiber, Paul Rosenberg

**Affiliations:** Department of Medicine, Duke University School of Medicine, Durham, NC

## Abstract

Cardiomyocytes in the sinoatrial node (SAN) are specialized to undergo spontaneous diastolic depolarization (DD) to create action potentials (AP) that serve as the origin of the heartbeat. Two cellular clocks govern DD: the membrane clock where ion channels contribute ionic conductance to create DD and the Ca^2+^ clock where rhythmic Ca^2+^ release from sarcoplasmic reticulum (SR) during diastole contributes pacemaking. How the membrane and Ca^2+^ clocks interact to synchronize and drive DD is not well understood. Here, we identified stromal interaction molecule 1 (STIM1), the activator of store operated Ca^2+^ entry (SOCE), in the P-cell cardiomyocytes of the SAN. Functional studies from STIM1 KO mice reveal dramatic changes in properties of AP and DD. Mechanistically, we show that STIM1 regulates the funny currents and HCN4 channels that are required to initiate DD and maintain sinus rhythm in mice. Taken together, our studies suggest that STIM1 acts as a sensor for both the Ca^2+^ and membrane clocks for mouse SAN for cardiac pacemaking.

## Introduction

In the heart, primary pacemaker cells that are located in the SAN give rise to an action potential (AP) during diastole that initiates sinus rhythm. APs from the SAN cardiomyocytes are readily distinguished from other cardiomyocytes by a characteristic diastolic depolarization (phase 4)^1^. Here, activation of the funny current (I_f_) during diastole triggers a series of ionic currents that contribute to the spontaneous APs. HCN4 channels are the molecular entity that underlies the pacemaker current (I_f_) and regulate the rate of firing of the SAN^2, 3^. HCN4 channels and related surface membrane channels establish the membrane clock that is needed for spontaneous rhythmic oscillation of membrane potential characteristic of pacemaking.

In addition to the membrane clock, it is now widely accepted that rhythmic release of Ca^2+^ stores from the sarcoplasmic reticulum (SR) during diastole is critically important for SAN automaticity. The Ca^2+^ clock hypothesis posits that local diastolic Ca^2+^ release from the internal stores in SANC activates Ca^2+^ extrusion via the Na^+^/Ca^2+^ exchanger on the sarcolemma, which, in turn provides an inward current for DD^4^. How internal Ca^2+^ stores in cardiomyocytes are maintained is not well understood, but requires Ca^2+^ pumps (SERCA2a) and store operated Ca^2+^ entry (Orai1)^5, 6^. Consistent with the emerging role of SOCE in excitable cells, we and others have recently reported that stromal interaction molecule 1 (STIM1) and SOCE are present in SANC and is important for Ca^2+^ signaling of primary pacemaking cells^5, 7, 8^. We have considered that STIM1-Ca^2+^ signaling might have an important role in coordinating the membrane clock and Ca^2+^ clock on a beat to beat basis for pacemaker activity. In the present study, we find that deletion of STIM1-SOCE from SAN cells markedly altered their action potentials which leads to substantial remodeling of additional currents, most notably I_f_. Using HCN4 current recordings from SANC and expressed in cultured cells, we show that STIM1 influences HCN4 channel gating directly. Our studies extend our understanding of the role for STIM1 in pacemaking to suggest that STIM1 is a link between the membrane clock and the Ca^2+^ clocks.

## Results

### STIM1 deletion and SAN action potential

Our previous studies established that STIM1 is enriched in the mouse SAN^5^. To better characterize the population of SANC expressing STIM1, we examined TEM images of heart sections taken from STIM1-reporter mice^9^. Here, β-galactosidase reaction product can be detected on TEM sections as a black precipitate (Figure 1A). Using this approach, STIM1-LacZ is detected in cardiomyocytes that resemble P-cells and transitional cells (Figure 1B), previously described in the mammalian SAN using TEM^10, 11^. P-cells are the conventional pacemaking cells in the SAN and are characterized by the lack of T-tubules, sparse myofibrils and numerous mitochondria. Notably, STIM1-LacZ is detected in the SR as it makes membrane contacts with the cell surface, mitochondrial outer membrane and the nuclear envelope (Figure1C). The STIM1-LacZ expression was specific to the P-cells and transitional cells as STIM1-LacZ and was not detected in the surrounding atrial cardiomyocytes or nerve fibers (Figure 1D).

**Figure 1:**
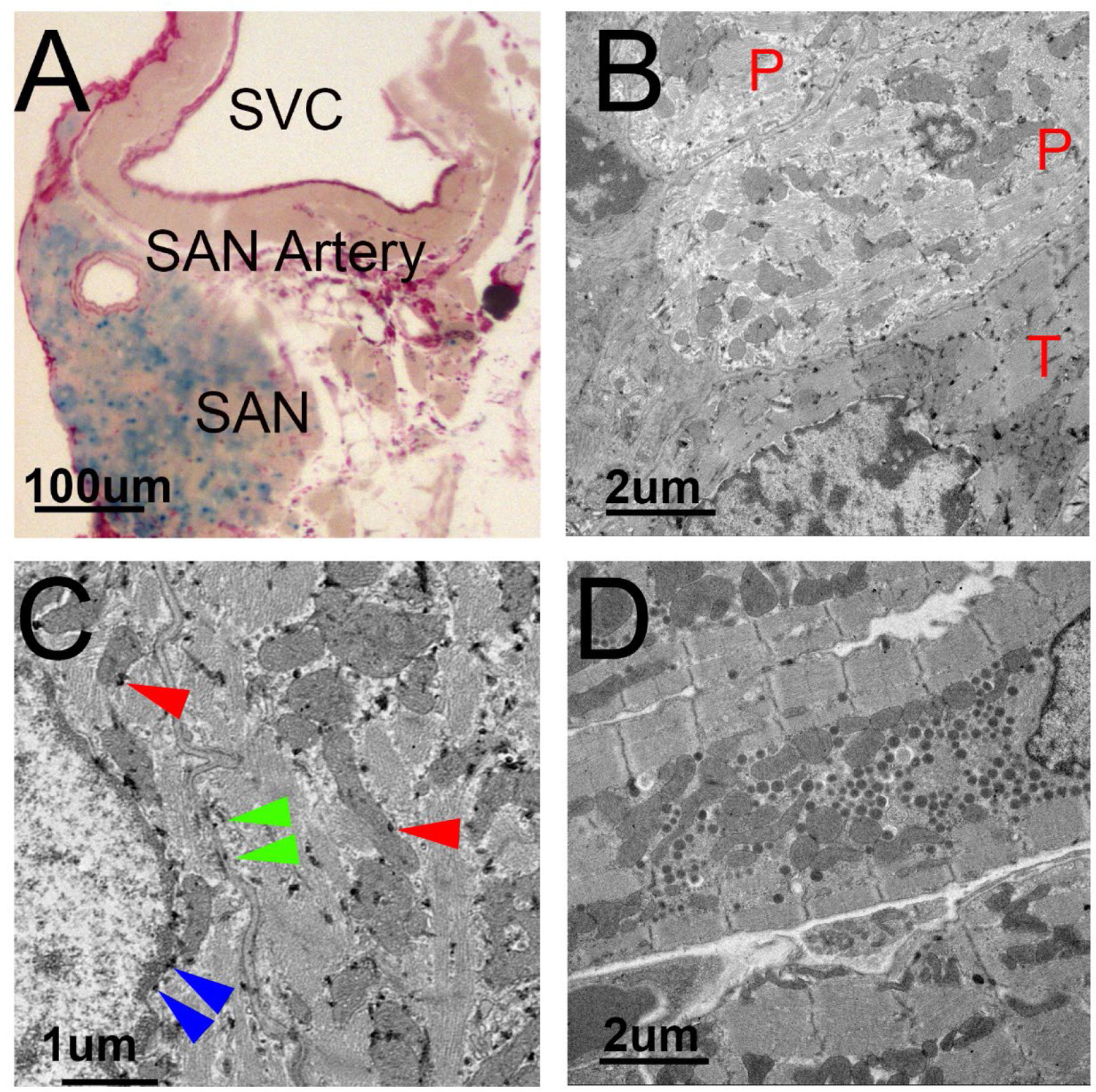
Localization of STIM1 in the mouse SANCs with TEM. (A) STIM1-LacZ staining in the mouse SAN (blue) and en bloc staining with uranyl acetate (pink). (B) STIM1 (dark precipitate) was present in a P cell (P) and transitional cell (T) in the mouse SAN with lower magnification. (C) STIM1 located in sub-plasma membrane (green arrows), mitochondria (red arrows) and sub-nuclear envelope (blue arrows). (D). STIM1 was not present in an adjacent atrial cell.

We have shown previously that cSTIM1^-/-^ mice exhibit reduced Ca^2+^ signaling, bradycardia and impaired HR response to hormonal stimulation^5^. To gain additional insight to the function of STIM1 in the SAN pacemaking, we measured spontaneous action potentials using microelectrode technique from SAN tissue preparations taken from STIM1^fl/fl^ (WT) and cSTIM1^-/-^ mice (KO). SA nodal cells (SANCs) were identified by specific morphologic features from brightfield microscopy as well as a slowly depolarizing action potentials consistent with DD recorded at 28°C (Figure 2, Table 1). Under current recording conditions, SAN firing rates were similar enabling us to compare AP properties without the confounding effects of heart rate variability. We observed that the maximum diastolic potential (MDP) was more negative in cSTIM1^-/-^ SAN cells compared to STIM1^fl/fl^ SAN cells (Figure 2A and Table 1). Because of the hyperpolarized MDP, the maximal excursion of the AP was greater in the cSTIM1^-/-^ SAN cells, as reflected in the larger AP amplitude (Table 1). Notably, we found an increase in the diastolic depolarization rate (DDR) in the SANs isolated from cSTIM1^-/-^ mice, implicating changes in the diastolic currents including I_f_ that influence AP properties (Figure 2B). These changes to APs from STIM1 deficient SANC led us to consider that adaptive changes to the membrane clock might compensate for altered Ca^2+^ signaling in cSTIM1^-/-^ mice. Of note, these changes to the APs were limited to pacemaker cells of the SAN as APs recorded from atrial tissue adjacent to the SAN were indistinguishable between STIM1^fl/fl^ and cSTIM1^-/-^ mice (Figure 2A and Table 1). Based on these changes, we proposed a working model where STIM1 activates Ca^2+^ entry (I_SOC_) during the repolarization and diastolic depolarization (AP phase 3 and 4) to replenish and maintain SR Ca^2+^ stores. As a result, the properties of the AP from SANC lacking STIM1 are markedly different.

**Figure 2:**
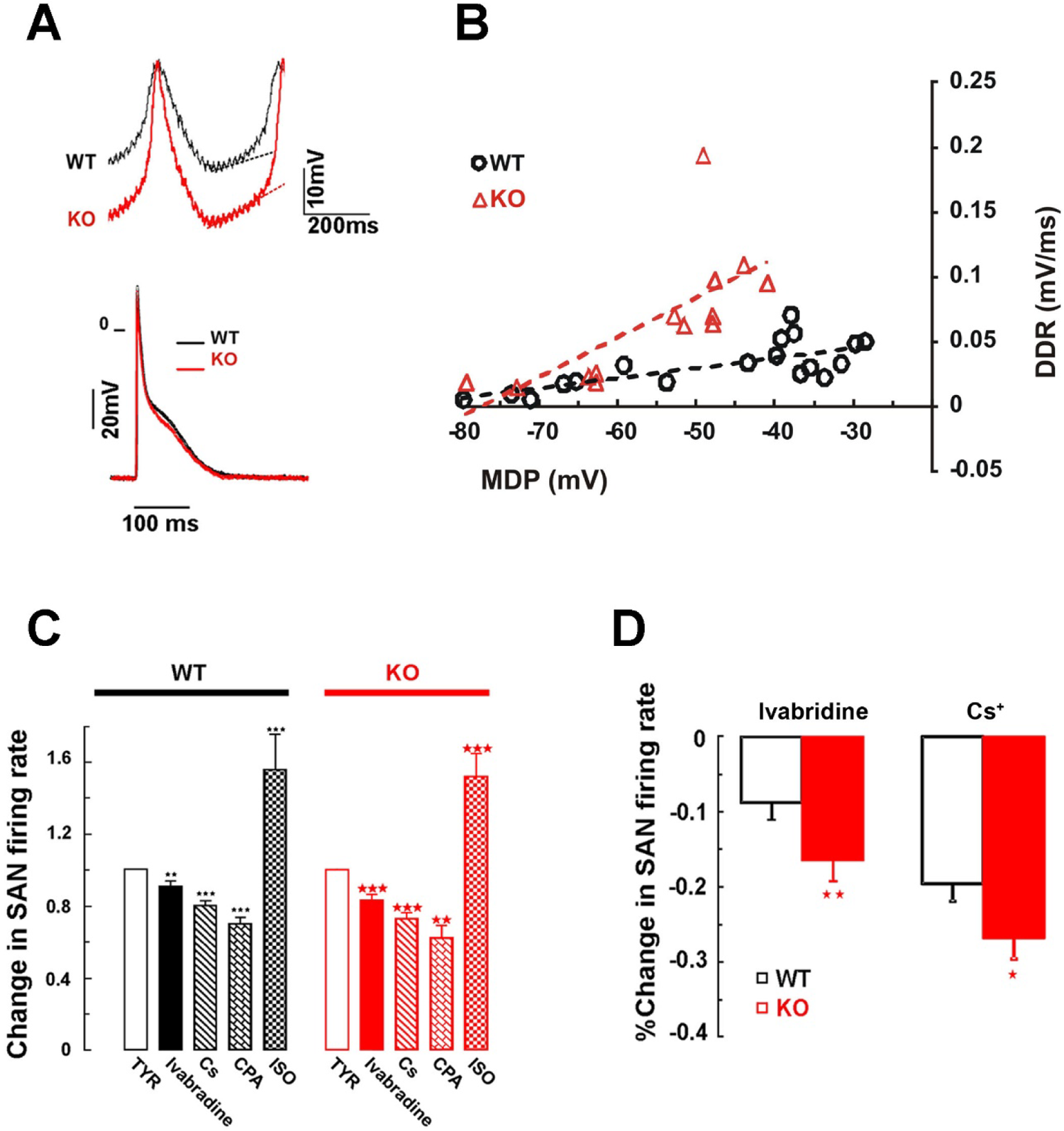
Deletion of STIM1 influences SAN action potential properties. (A) Intracellular recordings of action potentials of STIM1^fl/fl^ (WT, black) and cSTIM1^-/-^ (KO, red) SAN preparations (top) and atrial cells (bottom). (B) Diastolic depolarization rate (DDR) was plotted vs maximum diastolic potential (MDP). Each point represented an action potential recorded from STIM1^fl/fl^ SAN (open circle) and cSTIM1^-/-^ SAN (red triangle). No significant difference was noted in DDR when MDP was more negative than −60 mV. (C) Summary data for the WT (black column) and KO (red column) SAN firing rate change response to 10 µM Ivabradine (n=11), 2mM Cs^+^(n=15 for WT and n=9 for KO), 30 µM CPA (n=8 for WT and n=12 for KO) and 1µM isoproterenol (ISO, n=8 for WT and n=13 for KO). (D) Percentage change of STIM1^fl/fl^ and cSTIM1^-/-^ SAN firing rate in the absence and presence of ivbradine and Cs^+^. The significance was indicated by the asterisks (***p<0.001, **p<0.01 and *p<0.05)

**Table 1.**
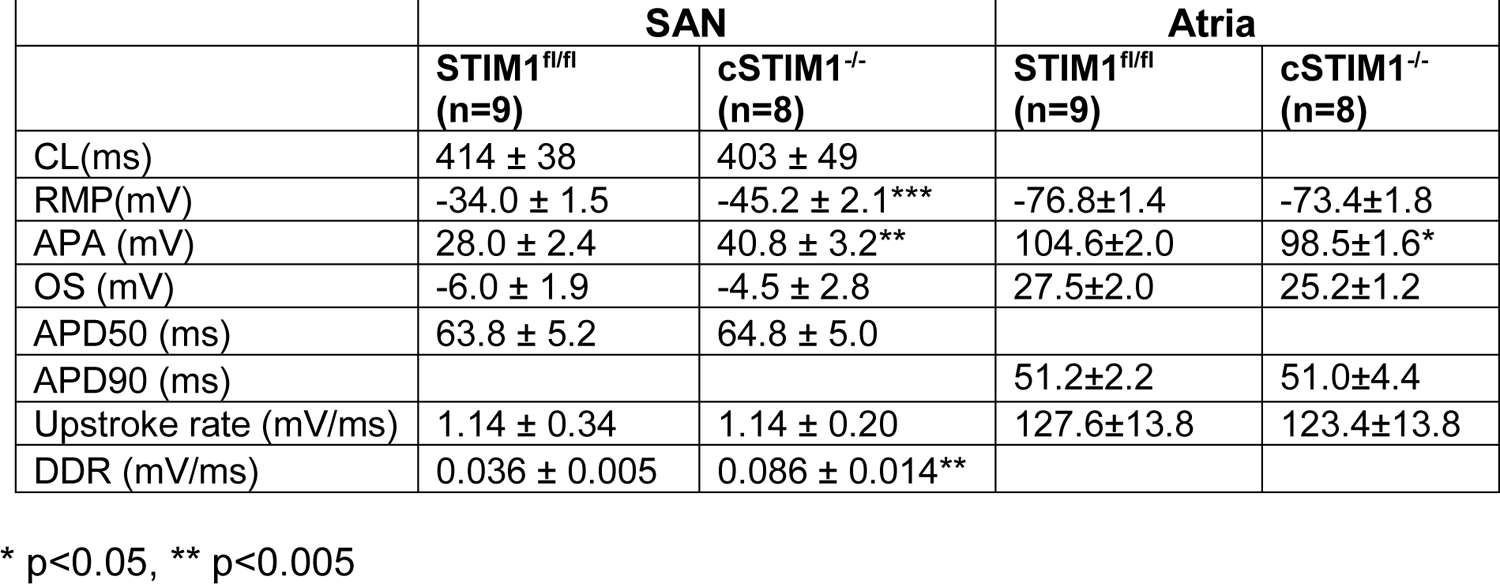
Parameters of action potentials of the STIM1^fl/fl^ and cSTIM1^-/-^ SAN

|  | **SAN** |  | **Atria** |  |
| --- | --- | --- | --- | --- |
|  | **STIM1<sup>fl/fl</sup>**<br>**(n=9)** | **cSTIM1<sup>-/-</sup>**<br>**(n=8)** | **STIM1<sup>fl/fl</sup>**<br>**(n=9)** | **cSTIM1<sup>-/-</sup>**<br>**(n=8)** |
| CL(ms) | 414 ± 38 | 403 ± 49 |  |  |
| RMP(mV) | -34.0 ± 1.5 | -45.2 ± 2.1*** | -76.8±1.4 | -73.4±1.8 |
| APA (mV) | 28.0 ± 2.4 | 40.8 ± 3.2** | 104.6±2.0 | 98.5±1.6* |
| OS (mV) | -6.0 ± 1.9 | -4.5 ± 2.8 | 27.5±2.0 | 25.2±1.2 |
| APD50 (ms) | 63.8 ± 5.2 | 64.8 ± 5.0 |  |  |
| APD90 (ms) |  |  | 51.2±2.2 | 51.0±4.4 |
| Upstroke rate (mV/ms) | 1.14 ± 0.34 | 1.14 ± 0.20 | 127.6±13.8 | 123.4±13.8 |
| DDR (mV/ms) | 0.036 ± 0.005 | 0.086 ± 0.014** |  |  |

\* p<0.05, \*\* p<0.005

To understand how the changes in the DD of the cSTIM1^-/-^ mice affected SAN function, we undertook pharmacologic studies. Cyclopiazoinc acid (CPA) is an inhibitor of the SR Ca^2+^ pump (SERCA2a) that results in SR Ca^2+^ depletion and has been a key reagent to probe the Ca^2+^ clock theory for pacemaking^4, 12^. An important consequence of CPA treatment is a reduction in the Ca^2+^ bound to STIM1 which eventually activates STIM1-dependent Ca^2+^ entry^7^. It is widely accepted that the mechanism by which CPA slows SAN firing is to deplete SR Ca^2+^ stores^13, 14^. Interestingly, CPA (30 µM) application to SAN from cSTIM1^-/-^ mice, which lacks I_SOC_, led to a dramatic slowing of the SAN firing (−52.3±9.9%, n=6 and p=0.035) compared to WT mice (−29.3±4.2%, n=5) These findings suggest that CPA acts not only to inhibit SERCA pumping but also to activate SOCE via STIM1 (Figure 2C and 2D).

We next used ivabradine (5 µM), an agent that slows SAN firing by specifically blocking HCN4 channels and alter DD, to assess the role of STIM1 in the regulation of the membrane clock^15^. We found that cSTIM1^-/-^ SANs were more sensitive to ivabradine as indicated by the reduction in SAN firing rate (8.8±2.2%, n=11) than STIM1^fl/fl^ mice (16.5±2.7%, n=11 and p=0.02 compared to WT) (Figure 2A and B). Similarly, SANs from cSTIM1^-/-^ mice exhibited more exaggerated slowing in response to Cesium (2 mM), strengthening the idea that HCN4 channels are sensitive STIM1. In contrast to the negative chronotropic effects of SERCA inhibition or HCN4 blockade, application of the β-adrenergic agonist isoproterenol accelerates SAN firing rates to the same level for cSTIM1^-/-^ and STIM1^fl/fl^ mice (Figure 2C). Isoproterenol increases cAMP production, which can influence HCN4 gating and Ca^2+^ cycling to increase rates of SAN firing^3^. Taken together, these studies demonstrate that STIM1 exerts complex actions on cardiac pacemaking by influencing not only the Ca^2+^ clock but might also modulate pacemaker currents I_f_ typically characterized in the membrane clock^5^.

To better define the relationship of STIM1 and the I_f_, we recorded hyperpolarization activated currents (I_f_) by whole cell patch-clamp technique from SAN cells isolated from STIM1^fl/fl^ and cSTIM1^-/-^ mice and found a four-fold increase in the I_f_ current density in the cSTIM1^-/-^ SANCs (Figure 3A and 3B). The change to I_f_ currents in cSTIM1^-/-^ SAN cells included a significant rightward shift in the fractional activation curve indicating that more HCN4 channels were activated at a given membrane potential compared to WT SAN cells (Figure 3C). To determine if changes to I_f_ were the direct result of STIM1, we infected cSTIM1^-/-^ SAN cells with an adenovirus carrying STIM1-cherry red to show I_f_ returned to control levels (supplemental Figure 1). Notably, STIM1-cherry partially co-localized intracellularly with HCN4 channels in SAN cells, as detected by immunofluorescent labeling for HCN4 (supplemental Figure 1D). Surprisingly, HCN4 channel expression was not different between WT and cSTIM1^-/-^ SANs, thus excluding greater HCN4 channel expression as the mechanism behind the augmented I_f_ currents in cSTIM1^-/-^ SAN cells (Figure 3D).

**Figure 3:**
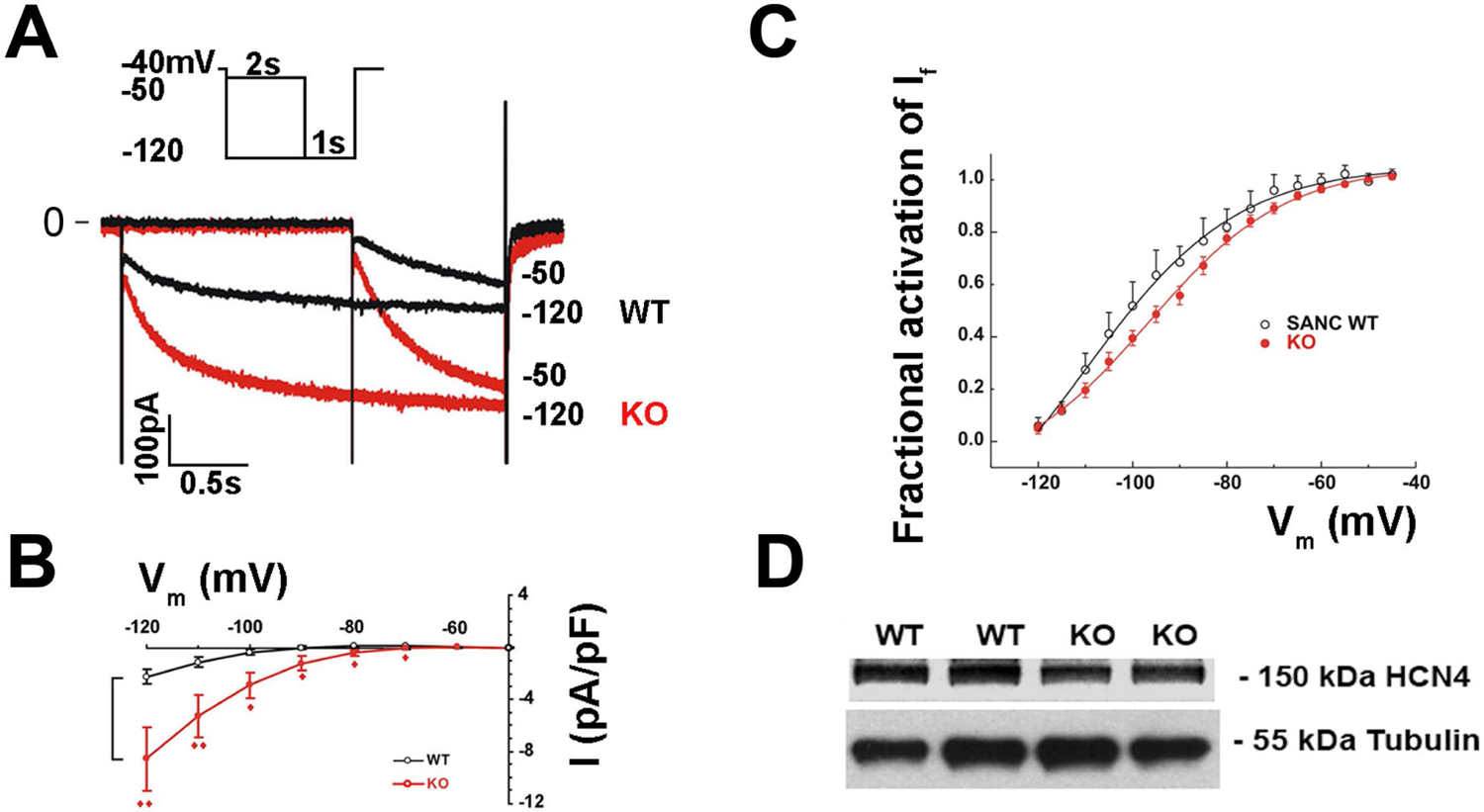
I_f_ currents recorded from STIM1^fl/fl^ (WT) and cSTIM1^-/-^ (KO) SAN cells. (A) Representative I_f_ currents recorded from STIM1^fl/fl^ (black) and cSTIM1^-/-^ (red) SAN cells. SAN cells were held at −40 mV and I_f_ currents were activated by hyperpolarization step from −50 to −120mV with an interval of 10mV. (B) I-V relationship of I_f_ currents are shown for STIM1^fl/fl^ (black, n=10) and cSTIM1^-/-^ (red, n=13) SAN cells. (C) Fractional activation curve of I_f_ was plotted by normalizing I_f_ currents activated during test pulse at −120mV and were averaged from STIM1^fl/fl^ (black, n=5) and cSTIM1^-/-^ (red, n=10) SANCs. Activation curve shifts 12 mV towards right (V_1/2_ from −108 to −96 mV) in the cSTIM1^-/-^ SANCs. (D) Western blot of HCN4 from two WT and KO SAN preparations.

We next considered that STIM1’s influence on the APs involves a direct interaction with HCN4 channels and therefore turned to a cell culture model where HCN4 channels are stably expressed in 293 cells. STIM1 overexpression in HCN4-293 cells eliminated the I_f_ current over a range of potentials tested (Figure 4A and 4B). In contrast, silencing endogenous STIM1 in HCN4-293 cells led to a dramatic increase in HCN4 current density (Figure 4A, B), consistent with the results from SAN cells of cSTIM1^-/-^ mice. In fact, we could demonstrate that HCN4 co-immunoprecipated with STIM1 using specific antibodies for STIM1 and HCN4 (Figure 4C). These studies provide a direct test of our idea that STIM1 regulates HCN4 currents and offers a plausible mechanism for the increased I_f_ in the SAN cells of cSTIM1^-/-^ mice. In Figure 2, an exaggerated slowing of the SAN firing rate was observed for cSTIM1^-/-^ SANs treated with CPA. Given that CPA-Ca^2+^ store depletion initiated STIM1 migration from the cortical ER to the cell membrane where it activates Orai channels, we reasoned that CPA might influence HCN4 current. Here, we show that CPA reduced HCN4 currents by 45% in HCN4 expressing HEK293 cells indicating that activation of STIM1 reduced HCN4 current (Figure 5A, B). Interestingly, immunostaining HEK293 cells with HCN4 specific antibodies revealed HCN4 channels were retained inside the cytoplasm for both control and 30 µM CPA treatment. However, a greater portion of HCN4 channel were clustered in a perinuclear compartment after CPA (Figure 5C). Together these studies support the notion that STIM1 and maintenance of Ca^2+^ stores are important for HCN4 trafficking and funny current activation.

**Figure 4:**
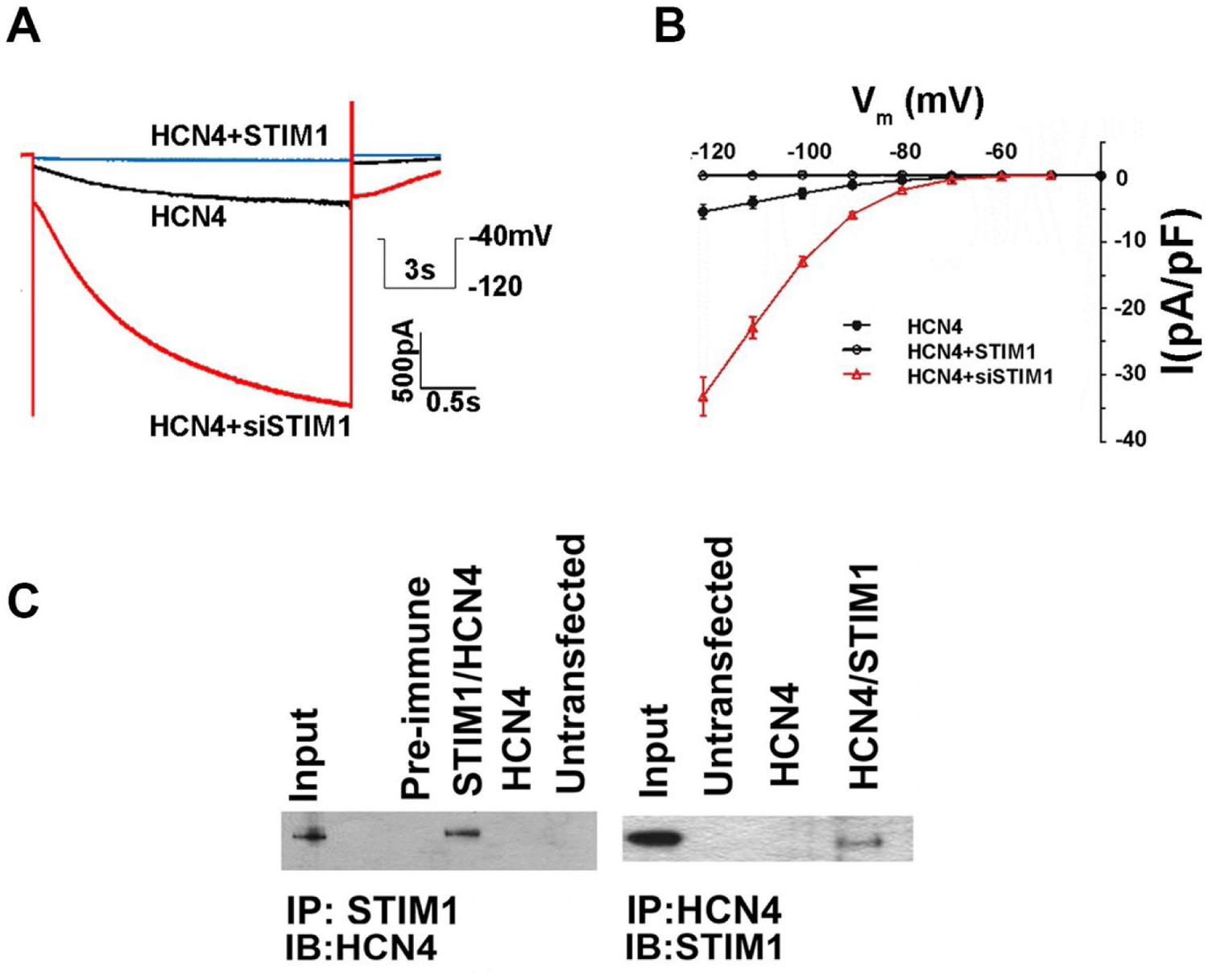
STIM1 physically interacts with HCN4 channels. (A) Representative current recordings for HCN4 currents stably expressed in HEK293 cells. Individual current recordings and I-V relations obtained from HEK-HCN4 cells transfected with plasmids encoding STIM1, scrambled shRNA, or shRNA for STIM1. (B) I-V relations for HCN4 currents in overexpressed STIM1 (open circle), silenced STIM1 (red, triangle) and control HCN4-HEK293 cells (filled circle, black). (C) Immunoprecipitation of HCN4 and STIM1 in HEK293 cells stably expressing HCN4. Lysates were incubated with antibodies for STIM1 (left) or HCN4 (right) and then immunoblotted for the HCN4 and STIM1 respectively. Input lysates were from HCN4/STIM1 expressing cells. Controls included untransfected or singly transfected cells.

**Figure 5:**
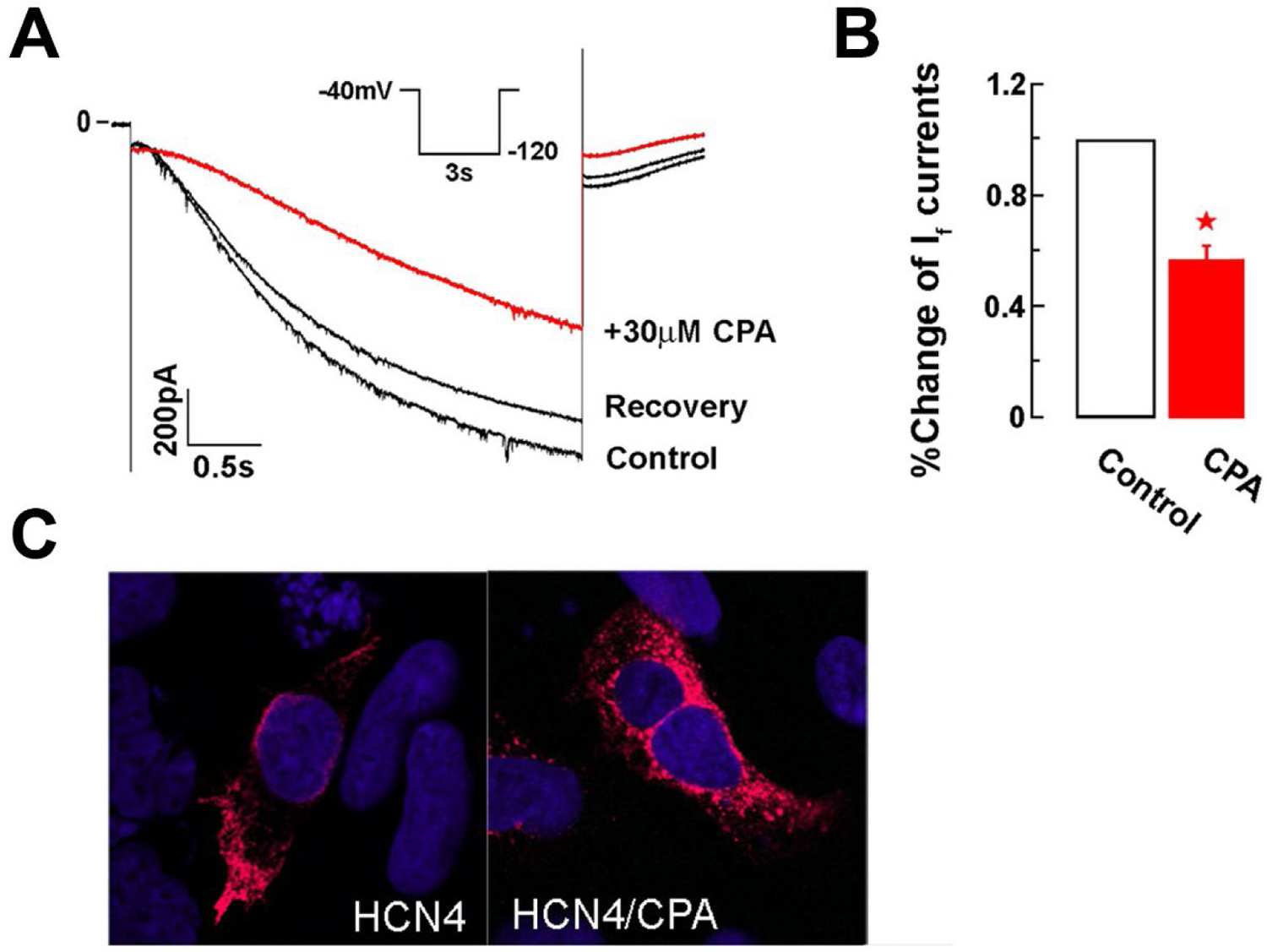
CPA-induced Ca^2+^ store depletion inhibit HCN4 current in stably HCN4 expressing HEK 293 cells. (A) Representative HCN4 currents recorded from HEK293 cells with stably expressing HCN4 channels. (B) Summarized data showed significant inhibition of HCN4 currents by external perfusion with 30µM CPA. (C) Immunostaining of HCN4 channels (red) in HEK293 cells in the absence and the presence of 30 µM CPA.

## Discussion

Automaticity, or the ability to self-generate action potentials, is an essential feature of the cardiac pacemaking cells of the SAN. At work in these specialized cardiomyocytes are ionic currents activated during the diastole to trigger the action potential and sinus rhythm. Our recent work showed that STIM1, a Ca^2+^ sensor in the sarcoplasmic reticulum, coordinated Ca^2+^ signaling in SAN cells and was needed for SAN function in the mouse^5^. In the present study, we extend our understanding of STIM1 in the SAN and suggest STIM1 links the Ca^2+^ and membrane clocks during pacemaking. Notably, action potentials recorded from cSTIM1^-/-^ SANs were markedly altered that suggested changes to early diastolic events, namely the MDP and DDR. Moreover, our investigation of the currents underlying DD identified a previously unrecognized relationship between STIM1 and HCN4 channels. Taken together these studies provide novel insight as to how STIM1 influences cardiac pacemaking that extends beyond Ca^2+^ signaling and directly involves components of the membrane clock.

Recently we reported that STIM1-SOCE has a key role in SAN cells where STIM1-Ca^2+^entry refills SR Ca^2+^ stores and thereby sustains the Ca^2+^ clock for cardiac pacemaking. We showed that store operated Ca^2+^ currents that act as inward background currents and counterbalance the repolarizing K currents typically influencing phase 3 of the AP. These K and SOC currents are active in the AP during late repolarization and early diastole phases and therefore influence the MDP that is a key determinant of DD^16–18^. Consistent with this role for SOCE during this phase, we found that APs from cSTIM1^-/-^ mice were markedly remodeled. Loss of the STIM1-Ca^2+^ entry in cSTIM1^-/-^ SAN led to unopposed K efflux during phase 3, which leads to dramatic changes to the MDP and DDR—key components of the DD. One consequence to the change in these AP properties of cSTIM1^-/-^ SAN was the upregulation of the HCN4 currents. Given the importance of HCN4 channels to DD and sinus node function, we explored the relationship of STIM1 and HCN4 channels. Our studies offer new and novel insight to cardiac pacemaking by introducing a new macromolecular complex that contributes to automaticity.

STIM1 directly influences many different ion channels, pumps and transporters including Orai1-3, Cav1.2, CLC, TRPC1, NCX1, SERCA1/2 and voltage-gated K channels (Kv1.3)^19–26^. Our findings, interpreted alongside these other studies, suggest that STIM1 in the SAN organizes HCN4 and Orai1 channels into local domains to facilitate cooperation between the membrane and Ca^2+^ clocks. We have recently shown that STIM1 is located in the ER membrane, near SERCA2a and phospholamban (PLN) of ventricular cardiomyocytes^27^. PLN can inhibit SERCA function and limit Ca^2+^ store refilling. STIM1 competes with SERCA for PLN and thereby can enhance SERCA function as was evident by the greater Ca^2+^ stores in the STIM1-expressing cardiomyocytes. Interestingly, both SERCA and PLN have important roles in SAN pacemaking and are key components of the Ca^2+^ clock^28^. Functionally, STIM1 deficient SAN cells exhibit marked reduction in the amplitude of Ca^2+^ signal and increased I_f_ from SAN cells. In the present work, we propose a model where STIM1 located in the SR connects to the sarcolemmal membrane in order to interact with HCN4 channels. We provide plausible ultrastructural evidence for the complex formation of SERCA/PLN, STIM1 and Orai1/HCN4. TEM studies localize STIM1-LacZ in the SR membrane adjacent the plasma membrane where ion channels reside to regulate DD. In this way, STIM1 coordinates refilling of Ca^2+^ stores and setting the MDP and rapid DDR that are critical for heart rate adaptability.

How does STIM1 influence I_f_? Certainly, the hyperpolarized MDP in the cSTIM1^-/-^ SAN provides the conditions to increase HCN4 channel gating and prevent further sinus slowing. We have considered that other mechanisms by which STIM1 might also influence the funny current. The signaling molecule cAMP binds to HCN4 channels and enhances gating of the channel^3^. Enzymes responsible for cAMP metabolism include adenylate cycles (AC) and phosphodiesterases and many of these enzymes are Ca^2+^ sensitive and directly respond to changes in SOCE and store depletion^29–32^. In this case, local changes in Ca^2+^ such as SOCE would enhance cAMP levels and augment HCN4 channel gating. However, in our studies SOCE and Ca^2+^ stores are reduced in SAN cells lacking STIM1. In this case, we would predict that AC and PPDE activity would be reduced in STIM1 depleted SANC resulting in reduced I_f_ current density. However, we found I_f_ density was much greater supporting the view that cAMP metabolism is not impaired. In addition, we note that isoproterenol, which increases cellular cAMP levels, increased SAN firing for the WT and cSTIM1^-/-^ mice to the same extent. In fact, we provide evidence to suggest STIM1 might influence the cellular trafficking of HCN4. STIM1 located in the ER might tether HCN4 intracellularly and thereby limiting its activation. Prior studies have focused on HCN4 channel assembly and show that HCN4 processing occurs in the ER^33^. In fact, disease-causing mutations in HCN4 have been identified in patients presenting with sinus node dysfunction and other cardiac phenotypes^34–36^. Maintaining the ER compartment would be important for biosynthesis and trafficking HCN4 channels and enable mature channels to insert into plasma membrane. STIM1 deficient cardiomyocytes have been shown to exhibit ER stress^37^. ER stress halts protein translation, upregulates chaperones to facilitate protein folding, and enables delivery of membrane proteins to the plasma membrane. It is entirely possible that STIM1 contributes to the processing and cycling of HCN4 channels to the cell membrane that governs HCN function^35^. When STIM1 was added back to cultured cSTIM1^-/-^ SAN myocytes, we found that HCN4 currents were rescued. STIM1 mediated channel processing in the ER is therefore a novel mechanism to control ion channel function.

In conclusion, we show that STIM1 located in SR of P-cells of the SAN that function to influence not only Ca^2+^ signaling, but also HCN4 channel activity. STIM1 and influence diastolic depolarization by remodeling Ca^2+^ entry and by affecting HCN4 activity. In this way, STIM1 acts to integrate the Ca^2+^ and membrane clocks. These findings introduce a novel mechanism to SAN automaticity and provide a mechanistic link between the membrane and Ca^2+^ clocks for SAN automaticity. A greater understanding of these mechanism will provide novel strategies to address sinus and atrial arrhythmias, which are major medical problem worldwide.

## Acknowledgements

We would like to thank Dr. Juliane Stieber (Univ. Erlangen) for graciously providing plasmid encoding human HCN4. This project was supported by Award Number R01HL093470 (PBR), from the National Institute of Health. The content is solely the responsibility of the authors and does not necessarily represent the official views of the National Institutes of Health.

## Materials and Methods

### Animals

STIM1-lacZ mice were genotyped as previously described^9^. STIM1^fl/fl^ mice (C57BL/6) were generated as previously described^38^. α-MHC (Cardiac specific)-cre transgenic mice (α-MHC-Cre) (C57BL/6) were obtained from Jackson Laboratories. To generate cardiac specific knockout of STIM1, αMHC-Cre transgenic mice were bred with founder STIM1^fl/fl^ mouse and the progeny STIM1^fl/-^;α-MHC-Cre^+/-^ were then back-crossed with STIM1^fl/fl^ mice. All mice were maintained in pathogen-free barrier facilities at Duke University and were used in accordance with protocols approved by the division of laboratory animal resources and institutional animal care & use committee at Duke University.

### Immunohistochemistry

For antibody staining, SAN cells or were isolated, plated in laminin coated 35mm Matek dish. SAN cells or HEK293 cells were then fixed in 4% paraformaldehyde for 10 minutes (for STIM1 and HCN4 staining). After fixation SAN cells were washed in PBS, blocked in normal goats serum and stained overnight in primary antibody at 4°C. STIM-1 polyclonal (Protein Tech group) was used at 1:250 and HCN4 polyclonal (Sigma) was used at 1:500. After washing, staining with the secondary antibody and counterstaining with DAPI slides were mounted in vectashield.

### Electron Microscopy and histology

For ultrastructural localization of STIM1-LacZ by TEM, hearts were fixed for 5 minutes in 2% PFA 0.2% Gluteraldehyde in PBS. SA nodes were then dissected and stained for LacZ as described in Stiber et al. ^9^. Tissue was post fixed in 2% PFA and 2% glutaraldehyde in 0.1 M phosphate buffer, pH 7.4. then fixed in 1% osmium tetraoxide, and stained en bloc with 1% uranyl acetate. Tissue was then was dehydrated in a graded ethanol series, taken through a series of Spurrs resin/Ethanol washes and embedded in Spurrs resin. Thick sections were cut to locate the SA node and then thin sections were cut at 70nm, mounted on copper grids and counterstained with 2% uranyl acetate and lead citrate .Thick sections were counterstained briefly on a hot plate with toluidine blue and basic fuschin in order to outline tissue but not mask LacZ staining.

### SAN preparation, SAN cell isolation and Adenoviral rescue

Procedures were adapted as previously described^39^. The heart was excised quickly from anaesthetized mouse and immersed into oxygenated Tyrode solution containing (in mM): NaCl 140, KCl 5.4, MgCl_2_ 1.05, NaH_2_PO_4_ 0.33, CaCl_2_ 1.8, HEPES 5, Glucose 10 and pH 7.4. For single SAN cell isolation, the SAN was dissected out from surrounding atrial tissues and cut into several small pieces. The SAN was then incubated in Ca^2+^-free solution containing (in mM): NaCl 140, KCl 5.4, MgSO_4_ 2, NaH_2_PO_4_ 0.33, HEPES 5, Glucose 10, Taurine 5 and pH 7.2, for 10 min at room temperature. The enzymatic digestion solution was prepared with Ca^2+^-free solution that contains BSA 0.2%, Collagenase (Type II, Worthington) 0.25 mg/ml, and Elastase 0.2 mg/ml. The digestion step (up to 45 min) was carried out in a shaking incubator at 37 °C, at a speed of 120 rpm. The single cells were released in modified KB solution by gentle titration with a glass transfer pipette. KB Solution includes: KCl 85, K Glutamate 20, KH_2_PO_4_ 20, Taurine 20, EGTA 0.5, Glucose 20, Creatine 5, Succinic Acid 5, Pyruvic Acid 5, MgSO_4_ 5, HEPES 5, pH 7.2. The isolated SAN cells were stored in KB solution at 4 °C at least for 2 hours before experiments. Those SAN cells having spindle or spider-like shape, beating in Tyrode solution, and showing I_f_ currents were chosen for patch clamp experiments.

For STIM1 rescue experiments, SAN cells were isolated from cSTIM1^-/-^ mice and cultured in MEM medium with L-glutamine (Mediatech). The additional supplements were added to medium including (in mM) 0.1% BAS, 2,3-butanedione monxime (BDM) 10, 1X Gibco insulin-selenium-transferrin, 1% penicillin-strptomycin, Creatine 5, taurine 5 and L-carnitine 2. The SANC were pre-plated in culture dish for 1 hr to remove fibroblast contamination. The SANCs were then transferred to a 35 mm petri dish and rSTIM1-cherry adenovirus with a multiplicity of infection (MOI) of 100 was added to culture medium. SANCs infected with β-gal were used as control. After 36-48 hr culture, SANC was plated to a perfusion chamber for I_f_ recordings.

### Electrophysiology

For intracellular action potential recording, the mouse SAN was pinned down to a Warner chamber with endocardium faced up and was then continuously perfused with oxygenated Tyrode solution, as previously described^5^. The action potentials of the SAN were recorded with conventional microelectrode technique. The microelectrode typically had a resistance of 20-60 MΩ and was filled 3M KCl. The primary SAN cells were identified by its specialized action potentials that show less negative diastolic potential, smooth transition from diastolic to upstroke and slower upstroke rate of action potentials.

For ionic current recording from single SAN cells, the SAN cells were placed in a Warner perfusion chamber, as previously described^5^. The patch pipette was pulled from borosilicate glass capillaries (Sutter Inc.) with a Sutter P-87 micropipette puller. The pipette usually had a resistance of 4-6 MΩ when filled pipette solution containing (in mM): Aspartate potassium 140, Mg·ATP 2, EGTA 11, ATP·Na_2_ 2, CaCl_2_ 1, HEPES 10 and pH 7.2. For I_f_ current recordings, perforated patch clamp technique was used. The final concentration of amphotericin B in pipette solution was 300 µg/ml. The liquid junction potential was nulled before gigiseal formation. Whole-cell currents were recorded using an AXON AXOPatch 200B amplifier connected to DIGIDATA 1322A digital converter. Whole cell currents were filtered at 2 KHz and sampled at a rate of 10 kHz. The data were collected through Clampex software 9.0 (Mol Device) and stored in computer for further analysis. All experiments were carried out at room temperature except those specified.

Plasmid DNA encoding HCN4 channels was a gift from Dr. Juliane Stieber (Univ. Erlangen). A HEK293 cell line stably expressing HCN4 channels was then established in this lab. For STIM1 and HCN4 interaction studies, STIM1 was transiently transfected into HEK293 cells with stable expression of HCN4 channels. After 48 hours culture in DMEM media containing 10% FBS, cells were harvested for patch clamp study or western blotting.

---

**Supplemental Figure 1:**
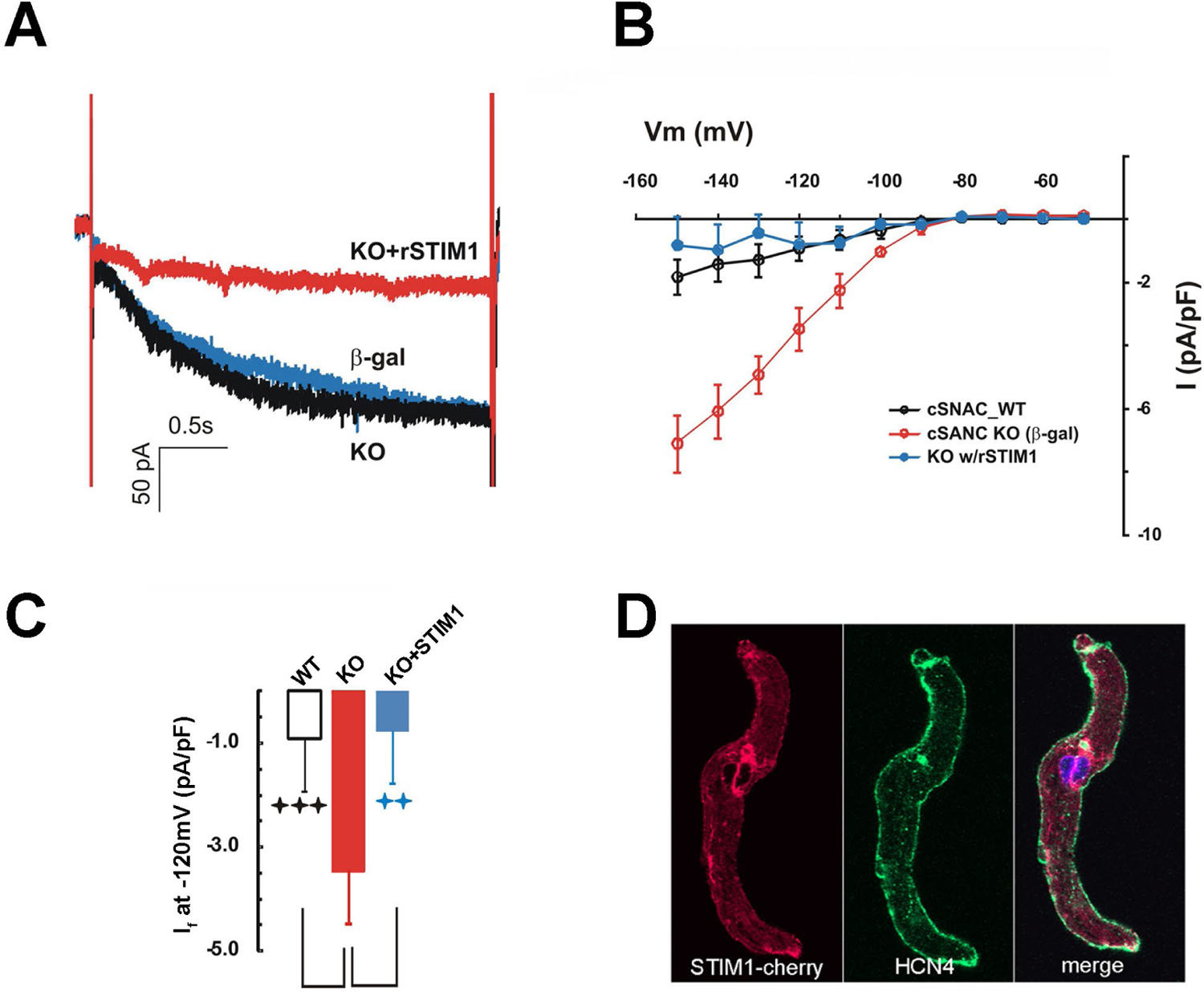
Rescue of I_f_ current in the cSTIM1^-/-^ SANCs. (A) Representative current recordings from cSTIM1^-/-^ SANC (black), cSTIM1^-/-^ SANCs infected with b-gal (blue) and with a STIM1-cherry adenovirus (red). The cultured SANCs were held −40mV and I_f_ currents were activated by a 3s hyperpolarization step from −50 to −150mV with an interval of −10mV. (B) I-V relationship of If currents in cultured cSTIM1^-/-^ SAN cells with (blue) or without (red) re-introduction of rat STIM1. (C) I_f_ current densities at −120mV from cultured STIM1^fl/fl^ (blank column, n=8, p<0.001 vs KO SANCs), cSTIM1^-/-^ (red, n=5) and cSTIM1^-/-^ SANC with rSTIM1 (n=7, p<0.01). (D) Immunostaining of HCN4 and overexpression of STIM1 m-cherry in cSTIM1^-/-^ SANC.

## References

1. Dobrzynski, H., Boyett, M.R. & Anderson, R.H. New insights into pacemaker activity: promoting understanding of sick sinus syndrome. Circulation 115, 1921–1932 (2007).

2. Stieber, J., et al. The hyperpolarization-activated channel HCN4 is required for the generation of pacemaker action potentials in the embryonic heart. Proc Natl Acad Sci U S A 100, 15235–15240 (2003).

3. Alig, J., et al. Control of heart rate by cAMP sensitivity of HCN channels. Proc Natl Acad Sci U S A 106, 12189–12194 (2009).

4. Bogdanov, K.Y., Vinogradova, T.M. & Lakatta, E.G. Sinoatrial nodal cell ryanodine receptor and Na(+)-Ca(2+) exchanger: molecular partners in pacemaker regulation. Circ Res 88, 1254–1258 (2001).

5. Zhang, H., et al. STIM1-Ca2+ signaling modulates automaticity of the mouse sinoatrial node. Proc Natl Acad Sci U S A 112, E5618–5627 (2015).

6. Vinogradova, T.M., et al. Sarcoplasmic reticulum Ca2+ pumping kinetics regulates timing of local Ca2+ releases and spontaneous beating rate of rabbit sinoatrial node pacemaker cells. Circ Res 107, 767–775.

7. Liou, J., et al. STIM is a Ca2+ sensor essential for Ca2+-store-depletion-triggered Ca2+ influx. Curr Biol 15, 1235–1241 (2005).

8. Roos, J., et al. STIM1, an essential and conserved component of store-operated Ca2+ channel function. J Cell Biol 169, 435–445 (2005).

9. Stiber, J., et al. STIM1 signalling controls store-operated calcium entry required for development and contractile function in skeletal muscle. Nat Cell Biol 10, 688–697 (2008).

10. Stoletzki, S., Schmiedl, A. & Richter, J. Intercalated clear cells or pale cells in the sinus node of canine hearts? An ultrastructural study. Anat Rec 260, 33–41 (2000).

11. Berger, J.M. & Rona, G. Fine structure of extra-nodal transitional cardiocytes in rat left atrium. J Mol Cell Cardiol 2, 181–185 (1971).

12. Maltsev, A.V., Yaniv, Y., Stern, M.D., Lakatta, E.G. & Maltsev, V.A. RyR-NCX-SERCA local cross-talk ensures pacemaker cell function at rest and during the fight-or-flight reflex. Circ Res 113, e94–e100 (2013).

13. Yaniv, Y., et al. New evidence for coupled clock regulation of the normal automaticity of sinoatrial nodal pacemaker cells: bradycardic effects of ivabradine are linked to suppression of intracellular Ca(2)(+) cycling. J Mol Cell Cardiol 62, 80–89 (2013).

14. Vinogradova, T.M., et al. Rhythmic ryanodine receptor Ca2+ releases during diastolic depolarization of sinoatrial pacemaker cells do not require membrane depolarization. Circ Res 94, 802–809 (2004).

15. DiFrancesco, D. The role of the funny current in pacemaker activity. Circ Res 106, 434–446 (2010).

16. Imtiaz, M.S., von der Weid, P.Y., Laver, D.R. & van Helden, D.F. SR Ca2+ store refill--a key factor in cardiac pacemaking. J Mol Cell Cardiol 49, 412–426.

17. Torrente, A.G., et al. Contribution of small conductance K(+) channels to sinoatrial node pacemaker activity: insights from atrial-specific Na(+) /Ca(2+) exchange knockout mice. J Physiol 595, 3847–3865 (2017).

18. Ho, W.K., Earm, Y.E., Lee, S.H., Brown, H.F. & Noble, D. Voltage- and time-dependent block of delayed rectifier K+ current in rabbit sino-atrial node cells by external Ca2+ and Mg2+. J Physiol 494 **(Pt** **3**), 727–742 (1996).

19. Liu, B., Peel, S.E., Fox, J. & Hall, I.P. Reverse mode Na+/Ca2+ exchange mediated by STIM1 contributes to Ca2+ influx in airway smooth muscle following agonist stimulation. Respir Res 11, 168.

20. Baryshnikov, S.G., Pulina, M.V., Zulian, A., Linde, C.I. & Golovina, V.A. Orai1, a critical component of store-operated Ca2+ entry, is functionally associated with Na+/Ca2+ exchanger and plasma membrane Ca2+ pump in proliferating human arterial myocytes. Am J Physiol Cell Physiol 297, C1103–1112 (2009).

21. Wang, Y., et al. The calcium store sensor, STIM1, reciprocally controls Orai and CaV1.2 channels. Science 330, 105-109.

22. Park, C.Y., Shcheglovitov, A. & Dolmetsch, R. The CRAC channel activator STIM1 binds and inhibits L-type voltage-gated calcium channels. Science 330, 101–105.

23. Prakriya, M., et al. Orai1 is an essential pore subunit of the CRAC channel. Nature 443, 230–233 (2006).

24. Huang, G.N., et al. STIM1 carboxyl-terminus activates native SOC, I(crac) and TRPC1 channels. Nat Cell Biol 8, 1003-1010 (2006).

25. Kunzelmann, K., et al. Role of the Ca2+-activated Cl-channels bestrophin and anoctamin in epithelial cells. Biol Chem 392, 125–134.

26. Feske, S., Prakriya, M., Rao, A. & Lewis, R.S. A severe defect in CRAC Ca2+ channel activation and altered K+ channel gating in T cells from immunodeficient patients. J Exp Med 202, 651–662 (2005).

27. Zhao, G., Li, T., Brochet, D.X., Rosenberg, P.B. & Lederer, W.J. STIM1 enhances SR Ca2+ content through binding phospholamban in rat ventricular myocytes. Proc Natl Acad Sci U S A 112, E4792–4801 (2015).

28. Li, Y., et al. CaMKII-dependent phosphorylation regulates basal cardiac pacemaker function via modulation of local Ca2+ releases. Am J Physiol Heart Circ Physiol 311, H532–544 (2016).

29. Lefkimmiatis, K., et al. Store-operated cyclic AMP signalling mediated by STIM1. Nat Cell Biol 11, 433–442 (2009).

30. Maiellaro, I., Lefkimmiatis, K., Moyer, M.P., Curci, S. & Hofer, A.M. Termination and activation of store-operated cyclic AMP production. J Cell Mol Med 16, 2715–2725 (2012).

31. Martin, A.C., et al. Capacitative Ca2+ entry via Orai1 and stromal interacting molecule 1 (STIM1) regulates adenylyl cyclase type 8. Mol Pharmacol 75, 830–842 (2009).

32. Putney, J.W., Jr. SOC: now also store-operated cyclase. Nat Cell Biol 11, 381–382 (2009).

33. Much, B., et al. Role of subunit heteromerization and N-linked glycosylation in the formation of functional hyperpolarization-activated cyclic nucleotide-gated channels. J Biol Chem 278, 43781–43786 (2003).

34. Nof, E., et al. Point mutation in the HCN4 cardiac ion channel pore affecting synthesis, trafficking, and functional expression is associated with familial asymptomatic sinus bradycardia. Circulation 116, 463–470 (2007).

35. Hardel, N., Harmel, N., Zolles, G., Fakler, B. & Klocker, N. Recycling endosomes supply cardiac pacemaker channels for regulated surface expression. Cardiovasc Res 79, 52–60 (2008).

36. Macri, V., et al. A novel trafficking-defective HCN4 mutation is associated with early-onset atrial fibrillation. Heart Rhythm 11, 1055–1062 (2014).

37. Collins, H.E., et al. Stromal interaction molecule 1 is essential for normal cardiac homeostasis through modulation of ER and mitochondrial function. Am J Physiol Heart Circ Physiol 306, H1231–1239 (2014).

38. Oh-Hora, M., et al. Dual functions for the endoplasmic reticulum calcium sensors STIM1 and STIM2 in T cell activation and tolerance. Nat Immunol 9, 432–443 (2008).

39. Hengtao Zhang, A.S., Jon Kim, Victoria Graham, Elizabeth A. Finch, Igor Nepliouev, Guiling Zhao, TianYu Li, Jonathan Lederer, Jonathan A. Stiber, Geoffrey S. Pitt, Nenad Bursac, Paul Rosenberg STIM1-Ca2+ signaling modulates automaticity of the mouse sinoatrial PNAS Accepted August 2015 (2015).

